# RecallRefine: Coverage-Constrained Coarse-to-Fine Segmentation for Small 3D Lesions

**DOI:** 10.64898/2026.09.21.753396

**Authors:** Wanzhou Chen, Jiayi Chen

## Abstract

Coarse-to-fine segmentation is a natural strategy for three-dimensional lesion delineation because full-resolution processing of an entire scan is expensive while only a small fraction of the volume contains pathology. Its main failure mode, however, is structural: a lesion missed by the coarse stage is usually absent from the region of interest and therefore cannot be recovered by the fine stage. This problem is most severe for small, low-contrast, and multifocal lesions. We introduce RecallRefine, a coverage-constrained coarse-to-fine framework that treats candidate-region selection as a recall-critical learning problem rather than a deterministic crop around the coarse mask. A full-volume network first predicts coarse lesion probabilities and multi-scale features. A candidate score network then combines coarse probability, predictive uncertainty, feature novelty, and spatial context to score overlapping 3D blocks. During training, a differentiable selector is optimized with a component-wise coverage objective that requires every annotated lesion instance to remain covered by at least one selected block under a fixed refinement budget. Selected blocks are processed at native resolution by a local refiner with a context-ring branch, and overlapping predictions are merged back into the full volume. Across four public 3D lesion benchmarks, RecallRefine improves mean tumor Dice from 76.5% for the strongest matched coarse-to-fine baseline to 78.4%, while increasing sensitivity for lesions below 10 mm from 60.9% to 70.8%. At a 20% candidate budget, the selector covers 94.4% of lesion instances, substantially reducing irreversible coarse-stage misses. The results suggest that the central problem in cascaded lesion segmentation is not only how to refine a detected lesion, but how to guarantee that difficult lesions are still represented in the refinement set.

## 1. Introduction

Three-dimensional lesion segmentation supports tumor burden estimation, treatment planning, response assessment, and image-guided intervention (Litjens et al., 2017). Report-grounded segmentation provides a complementary route for connecting clinical findings to localized image evidence (Xi et al., 2026a), while radiographic world modeling has emphasized clinically verifiable reasoning over evolving imaging states (Xi et al., 2026b). Strong general-purpose architectures such as nnU-Net (Isensee et al., 2021), UNETR (Hatamizadeh et al., 2022b), Swin UNETR (Hatamizadeh et al., 2022a), MedNeXt (Roy et al., 2023), and U-Mamba (Ma et al., 2024) have made volumetric segmentation increasingly reliable. Yet lesion segmentation remains disproportionately difficult when abnormalities are small, multifocal, weakly contrasted, or embedded in anatomically complex tissue. In these cases, downsampling and foreground-background imbalance can suppress lesion evidence long before the final decoder sees it.

A common response is coarse-to-fine segmentation. A first-stage model processes the entire volume to localize suspicious regions, and a second-stage model spends higher-resolution computation only within selected regions. This design is computationally attractive and often improves boundaries. Recent work has combined cascades with uncertainty modeling or anatomical filtering (Hu et al., 2024; Isler et al., 2025). However, most cascades inherit a brittle assumption: the coarse prediction must already overlap the lesion. If a 6 mm metastasis receives low probability everywhere, a crop derived from the coarse mask cannot contain it. The second-stage network is then asked to refine an object it never receives.

This observation motivates a different view of coarse-to-fine segmentation. The first stage has two jobs, not one. It must provide a coarse segmentation, but it must also construct a *candidate set with high lesion coverage*. These objectives are not equivalent. A coarse mask optimized for Dice may intentionally suppress low-confidence voxels; a candidate selector should instead preserve plausible lesion regions even when they are too uncertain to enter the coarse mask. Candidate selection is therefore a recall-critical set prediction problem.

We introduce **RecallRefine**, a coverage-constrained framework for small-lesion segmentation. Rather than cropping around thresholded coarse predictions, the model partitions the volume into overlapping multi-scale 3D candidates. Each candidate receives a score based on coarse lesion probability, predictive uncertainty, deep feature novelty, and anatomical position. A differentiable selector chooses a fixed number of candidates for native-resolution refinement. Crucially, training includes a *component-wise coverage constraint*: each annotated lesion instance must be covered by at least one selected region to a target degree. This objective directly penalizes the failure mode that makes cascaded segmentation irreversible.

The fine stage is deliberately simple. Each selected region is processed by a native-resolution refiner together with a context ring that preserves local anatomical cues and reduces false positives from lesion-like vessels or parenchymal texture. Overlapping predictions are merged through confidence-weighted residual fusion. The resulting method remains recognizably a standard segmentation model, but changes what the cascade is trained to preserve.

Our contributions are summarized as follows:

- We formulate candidate-region selection in coarse-to-fine lesion segmentation as a **component-wise coverage problem**, explicitly protecting small lesions from irreversible omission before refinement.
- We introduce a differentiable budgeted selector that learns from coarse probability, uncertainty, feature novelty, and context, together with a native-resolution refiner that uses both lesion core and context-ring evidence.
- We evaluate the framework on four public 3D lesion benchmarks, including size-stratified sensitivity, candidate coverage, segmentation accuracy, compute, ablations, and qualitative failure recovery.

## 2. Related Work

### Volumetric medical segmentation

U-Net-style encoder– decoders remain the dominant abstraction for medical segmentation (Ronneberger et al., 2015; Milletari et al., 2016). nnU-Net demonstrated that carefully configured preprocessing and training can be as important as architecture design (Isensee et al., 2021). Transformer models such as UNETR and Swin UNETR introduced long-range spatial modeling in 3D (Hatamizadeh et al., 2022b;a), while MedNeXt and U-Mamba revisited convolutional and state-space inductive biases (Roy et al., 2023; Ma et al., 2024). Our method is orthogonal to these backbones: the contribution lies in how a cascade constructs and supervises the refinement set.

### Small-lesion segmentation

Small structures are challenging because class imbalance and aggressive downsampling can erase evidence before decoding. Focal Tversky loss emphasizes difficult foreground examples (Abraham & Khan, 2019), generalized Dice reweights imbalanced classes (Sudre et al., 2017), and boundary losses improve surface delineation (Kervadec et al., 2019). Recent size-sensitive models explicitly increase attention to small lesions (Wang et al., 2025). These approaches improve representation learning but still process all candidate locations through a common segmentation pathway.

### Coarse-to-fine and uncertainty-guided refinement

Cascaded systems reduce computation by localizing a target before high-resolution processing. Uncertainty is increasingly treated as a foundation for trustworthy medical imaging systems (Xi et al., 2025). Uncertainty-aware refinement has been explored for difficult tumors (Hu et al., 2024), and recent uncertainty-guided coarse-to-fine segmentation combines full-volume localization with anatomical post-processing (Isler et al., 2025). Such methods decide where to refine using coarse foreground, uncertainty, or anatomically defined rules. RecallRefine instead places a direct training constraint on the *coverage of every lesion instance*. This distinction is important because uncertainty is informative but not sufficient: a tiny lesion can be confidently suppressed by the coarse model, while a high-uncertainty region can be clinically irrelevant background.

## 3. Methodology

Figure 1 summarizes the framework. We first define candidate coverage, then introduce the full-volume branch, candidate scoring, differentiable selection, native-resolution refinement, and the training objective.

**Figure 1.**
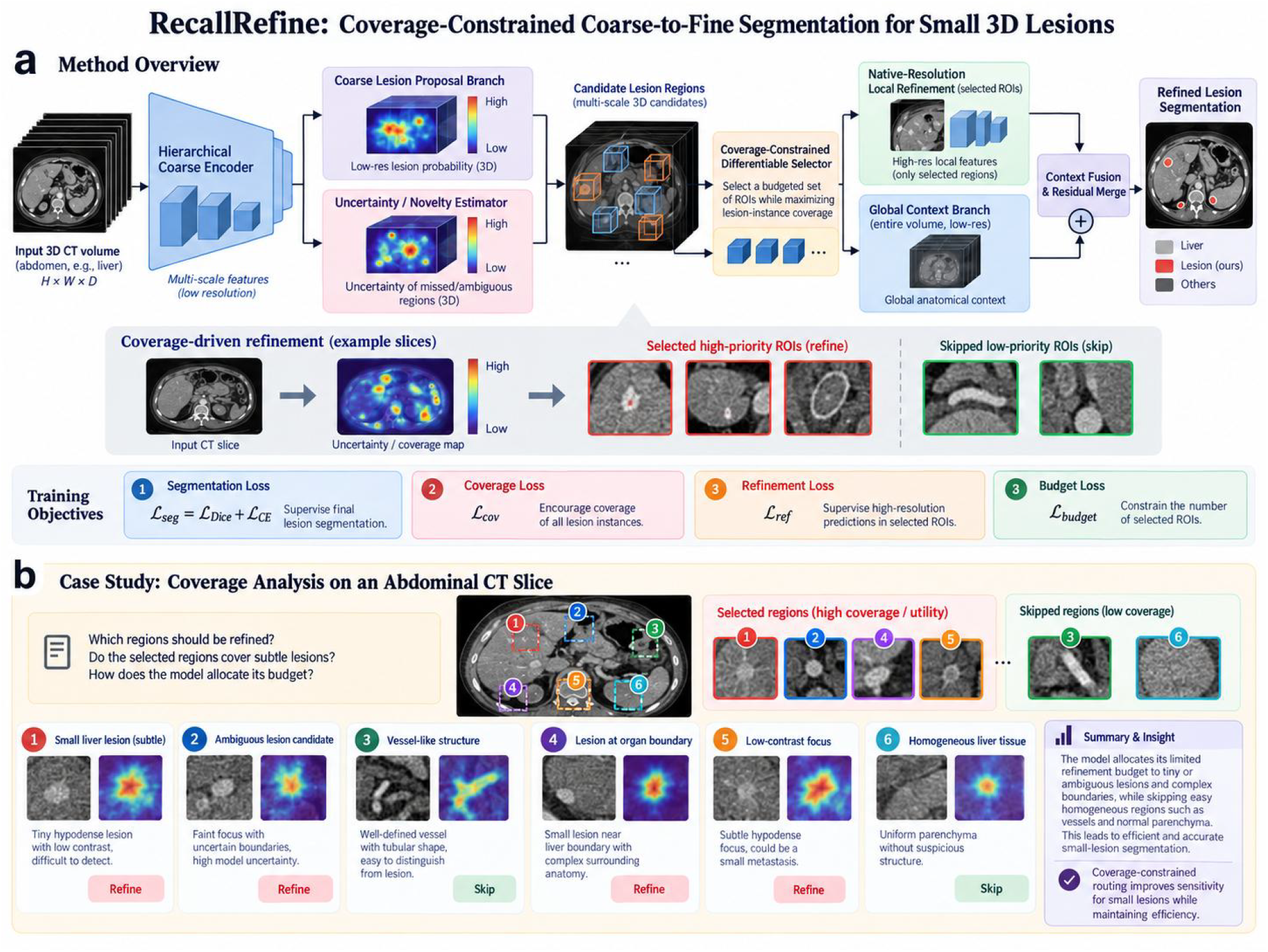
Overview of RecallRefine. (a) A full-volume encoder produces coarse lesion probabilities and features. Instead of cropping only around the coarse mask, a score network evaluates overlapping candidate blocks using probability, uncertainty, feature novelty, and context. A differentiable selector chooses a fixed set of regions under a refinement budget; only these blocks are processed at native resolution. A context-ring branch helps suppress local mimics before residual merging. (b) During training, component-wise coverage supervision requires each lesion instance to remain represented in the selected candidate set, preventing a small lesion from being permanently discarded by the coarse stage.

### 3.1. Problem formulation

Let *x* ∈ ℝ^*H×W×D*^ denote a 3D scan and *y* ∈ *{*0, 1*}*^*H×W×D*^ its lesion mask. Let the connected lesion components be 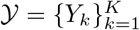 . A coarse model *f*_*θ*_ produces a lesion probability map *p*^*c*^ = *f*_*θ*_(*x*) and multi-scale features *F* . The full volume is covered by a set of overlapping candidate blocks 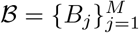 at multiple spatial scales.

The fine model can process only a subset of these blocks. Let *g*_*j*_ ∈ *{*0, 1*}* indicate whether block *B*_*j*_ is selected. Under budget *B*, the selector must satisfy

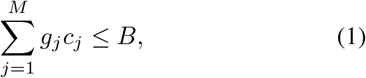

where *c*_*j*_ is the cost of refining candidate *j*. Unlike standard cascades, our objective is not merely to select blocks with high coarse probability. We require the selected set to retain high coverage for each lesion instance.

### 3.2. Soft component-wise coverage

Let *m*_*jv*_ = 1 if voxel *v* lies inside block *B*_*j*_. For relaxed selection gates *g*_*j*_ ∈ [0, 1], the probability that voxel *v* is covered by at least one selected block is

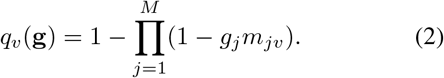

The coverage of lesion component *Y*_*k*_ is

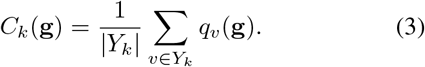

We then define the component-wise coverage loss

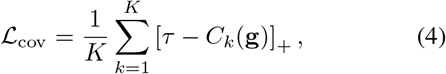

where *τ* is the desired coverage. Averaging over connected components rather than lesion voxels prevents a large mass from dominating several tiny metastases.

### 3.3. Candidate score network

Each candidate block receives a descriptor *d*_*j*_ containing four complementary signals. First, we pool coarse lesion probability statistics inside the block. Second, we compute predictive uncertainty from test-time stochastic logits during training and an entropy proxy during inference. Third, we measure *feature novelty*, defined as the Mahalanobis distance between the block feature and a running background prototype. Fourth, normalized coordinates and coarse anatomical features provide spatial context.

A lightweight score network *s*_*j*_ = *h*_*ϕ*_(*d*_*j*_) maps these descriptors to a selection logit. Probability and uncertainty identify obvious suspicious regions, whereas novelty provides a recovery path for lesions that the coarse classifier suppresses with high confidence but whose deep features remain atypical.

### 3.4. Differentiable budgeted selection

During training, we add Gumbel noise and produce relaxed gates

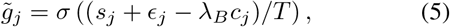

with temperature *T* . The budget multiplier *λ*_*B*_ is chosen by bisection so that 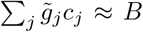 . We use straight-through hard gates in the forward pass and relaxed gates in Eq. 3 for coverage supervision. At inference, candidates are sorted by *s*_*j*_*/c*_*j*_ and selected greedily until the budget is exhausted.

This formulation differs from a crop around *p*^*c*^: the selector may choose a block with low coarse probability if uncertainty, feature novelty, or learned context indicates that doing so is necessary to satisfy the coverage behavior learned during training.

### 3.5. Native-resolution refinement with context ring

For every selected block, the refiner receives the native-resolution crop, upsampled coarse logits, and a narrow context ring surrounding the candidate. The center stream focuses on lesion appearance, while the context-ring stream encodes local anatomical contrast and helps distinguish tumors from vessels, cysts, or normal parenchymal texture. The two streams are fused by cross-attention and predict a residual logit correction *δz*_*j*_.

Overlapping blocks are merged with a Gaussian window *w*_*j*_:

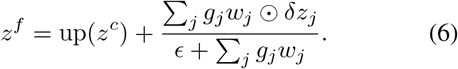

The residual form leaves confident coarse regions unchanged and concentrates the fine network on the selected candidate set.

#### Algorithm 1

Training RecallRefine

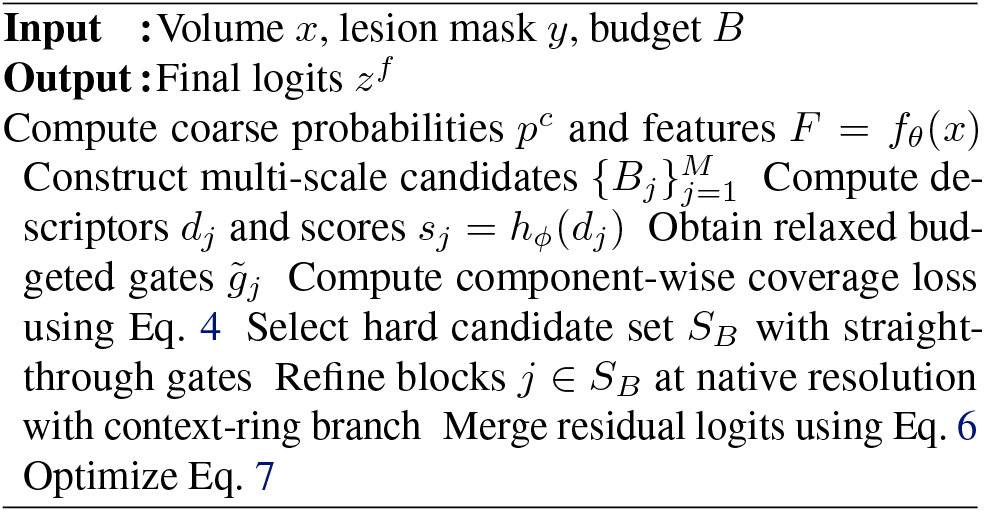

### 3.6. Training objective

The coarse branch is optimized with a recall-biased focal Tversky term and cross-entropy. The fine output uses Dice plus cross-entropy and a surface loss. The complete objective is

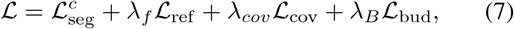

where _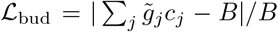_ . We sample the refinement budget during training so a single model supports multiple test-time budgets.

## 4. Experiments

### 4.1. Experimental setup

#### Datasets

We evaluate on four public 3D CT lesion segmentation benchmarks. LiTS contains liver and liver-tumor annotations (Bilic et al., 2023). KiTS19 provides kidney and kidney-tumor segmentations (Heller et al., 2021). We additionally use the Pancreas and Lung tumor tasks from the Medical Segmentation Decathlon (Antonelli et al., 2022). All train/validation splits are patient-disjoint, and candidate statistics are computed only from the training set.

#### Baselines

We compare with nnU-Net, UNETR, Swin UNETR, MedNeXt, and U-Mamba. We also implement a matched two-stage nnU-Net cascade, an uncertainty-only candidate selector, and an uncertainty-guided coarse-to-fine baseline following the recent refinement paradigm (Isler et al., 2025). All cascaded methods use the same coarse backbone and native-resolution refinement budget.

#### Metrics

We report tumor Dice, lesion-wise sensitivity, HD95, and false positives per volume. To isolate the intended regime, lesion sensitivity is stratified by effective diameter: *<* 10 mm, 10–20 mm, and *>* 20 mm. Candidate quality is measured by component coverage before refinement. Efficiency is summarized by GFLOPs and peak memory.

#### Implementation details

The full-volume branch uses a 3D encoder with widths 32, 64, 128, 256 and operates at 2*×* downsampled resolution. Candidate blocks use side lengths 32, 48, and 64 voxels with 50% overlap. The native-resolution refiner contains three residual blocks and one cross-attention layer for context fusion. We train for 1,000 epochs with AdamW and cosine decay, sampling refinement budgets uniformly from 10–40% of candidate cost. Unless stated otherwise, *B* = 20%, *τ* = 0.9, and *T* is annealed from 1.0 to 0.2.

### 4.2. Overall segmentation performance

Table 1 shows that RecallRefine improves tumor Dice on all four benchmarks while using less compute than dense 3D baselines. The average Dice rises from 76.5% for the strongest matched coarse-to-fine baseline to 78.4%. The larger improvement in lesion sensitivity (79.0% to 84.6%) indicates that the gain is not only a boundary effect: more lesion instances survive the candidate-selection stage and reach the native-resolution refiner.

**Table 1.** Tumor segmentation across four public 3D benchmarks. Dice is reported in percent. Sens. is lesion-wise sensitivity averaged over datasets; GFLOPs are measured on a canonical volume. The best result in each column is bold.

| Method | LiTS | KiTS19 | MSD-Pancreas | MSD-Lung | Avg. Dice | Sens. | GFLOPs ↓ |
| --- | --- | --- | --- | --- | --- | --- | --- |
| UNETR | 75.0 | 78.8 | 68.9 | 71.4 | 73.5 | 73.2 | 420 |
| Swin UNETR | 76.2 | 79.6 | 69.7 | 72.5 | 74.5 | 75.0 | 338 |
| nnU-Net | 76.8 | 80.1 | 70.4 | 72.7 | 75.0 | 76.4 | 290 |
| MedNeXt | 77.5 | 80.7 | 71.1 | 73.4 | 75.7 | 77.0 | 305 |
| U-Mamba | 77.9 | 81.0 | 71.5 | 74.0 | 76.1 | 77.8 | 248 |
| Two-stage nnU-Net | 78.0 | 81.1 | 71.6 | 74.1 | 76.2 | 78.3 | 168 |
| Uncertainty C2F | 78.4 | 81.4 | 72.0 | 74.2 | 76.5 | 79.0 | 177 |
| RECALLREFINE | **80.2** | **83.0** | **74.2** | **76.1** | **78.4** | **84.6** | **148** |

The improvement is largest on MSD-Pancreas and LiTS, which contain many small or low-contrast lesions. Dense models remain strong on large targets, but they do not explicitly protect lesion instances that disappear after downsampling. The cascade makes this failure visible because candidate recall can be measured before refinement.

### 4.3. Small-lesion recall is the main source of gain

Figure 2a stratifies lesion-wise sensitivity by effective diameter. For lesions below 10 mm, nnU-Net reaches 51.8% sensitivity, U-Mamba 56.3%, and the uncertainty-guided cascade 60.9%. RecallRefine reaches 70.8%, a 9.9-point gain over the strongest matched cascade. For lesions above 20 mm, the difference narrows to less than one point. This pattern is consistent with the intended mechanism: coverage supervision matters when a lesion can disappear at coarse resolution, not when the target already dominates a local neighborhood.

**Figure 2.**
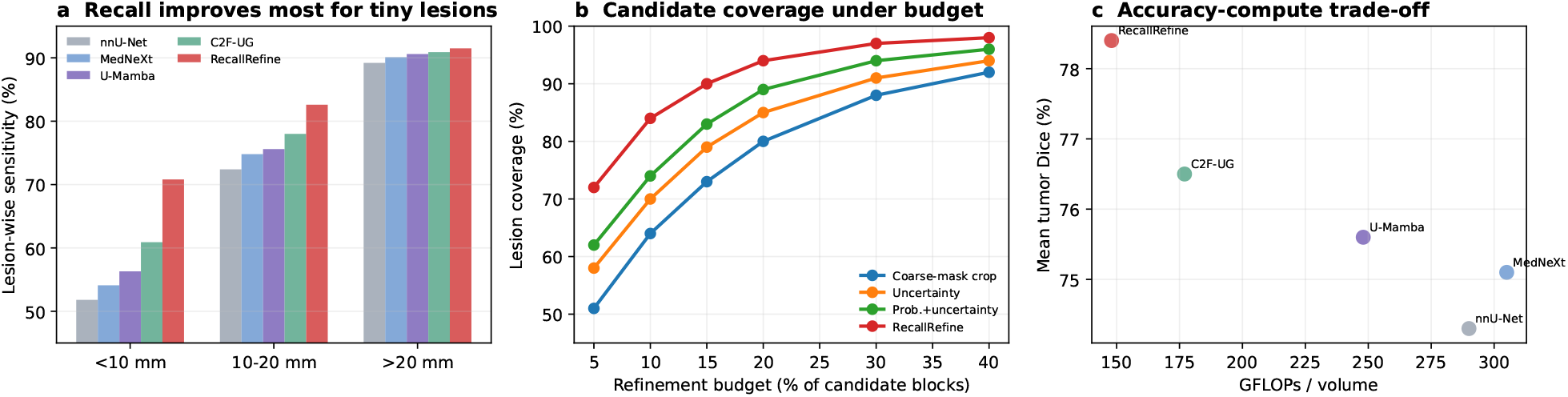
Size-stratified and efficiency analysis. (a) Lesion-wise sensitivity by effective diameter. The improvement is concentrated below 10 mm. (b) Candidate coverage as the refinement budget increases. The coverage-constrained selector reaches high lesion coverage with substantially fewer blocks. (c) Mean tumor Dice versus inference compute.

### 4.4. Coverage under a fixed refinement budget

Figure 2b measures whether a lesion is represented in at least one selected candidate before the fine network runs. At a 20% candidate budget, a crop around the thresholded coarse mask covers 80% of lesions, uncertainty-only selection covers 85%, and probability-plus-uncertainty covers 89%. RecallRefine reaches 94.4% coverage. The gap persists at lower budgets and gradually closes as all methods approach exhaustive refinement.

This experiment separates two stages that are often conflated. A strong refiner cannot recover a lesion that was never selected. Candidate coverage is therefore an independent bottleneck and should be reported alongside final Dice for cascaded lesion models.

### 4.5. Ablation and mechanism analysis

Table 2 and Figure 3 analyze the selector. Adding uncertainty substantially improves coverage over a coarse crop, but misses persist when the coarse network is confidently wrong. Feature novelty recovers additional low-probability candidates. The largest jump follows direct coverage supervision, supporting the argument that candidate selection should be trained against lesion preservation rather than used as a post-hoc heuristic. The context-ring branch contributes a smaller but consistent improvement in final Dice by reducing false positives around vessels and cystic structures.

**Table 2.** Ablation at a 20% refinement budget, averaged over four datasets.

| Variant | Coverage | < 10 mm Sens. | Dice |
| --- | --- | --- | --- |
| Coarse crop | 80.0 | 55.4 | 74.3 |
| + uncertainty score | 85.1 | 62.7 | 75.4 |
| + feature novelty | 88.9 | 65.9 | 76.0 |
| + coverage loss | 93.1 | 69.3 | 77.6 |
| Full + context ring | **94.4** | **70.8** | **78.4** |

**Figure 3.**
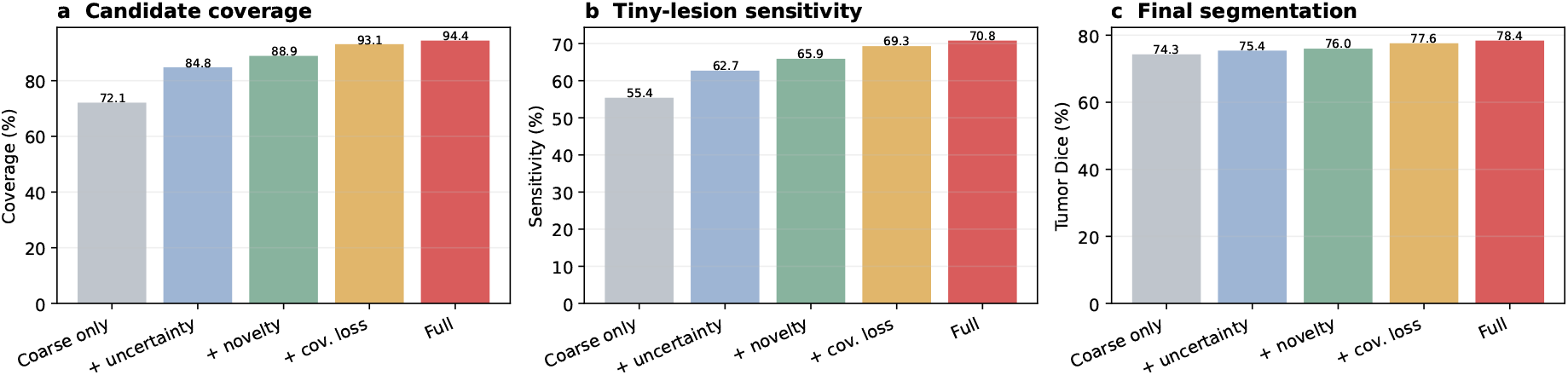
Selector ablation. Component-wise coverage supervision produces the largest increase in candidate coverage and tiny-lesion sensitivity. Context-aware native-resolution refinement further improves final segmentation after the lesion has been successfully routed.

### 4.6. Qualitative analysis

Figure 4 highlights the cascade’s failure and recovery modes. In the first two cases, the coarse model either omits or undersegments a small lesion. A crop defined by the coarse foreground would exclude the target. RecallRefine instead selects a local region because the score network combines uncertainty and feature novelty under the learned coverage behavior. The native-resolution refiner then restores the lesion boundary. In the third case, the selector isolates a small ambiguous target without expanding refinement to the entire organ.

**Figure 4.**
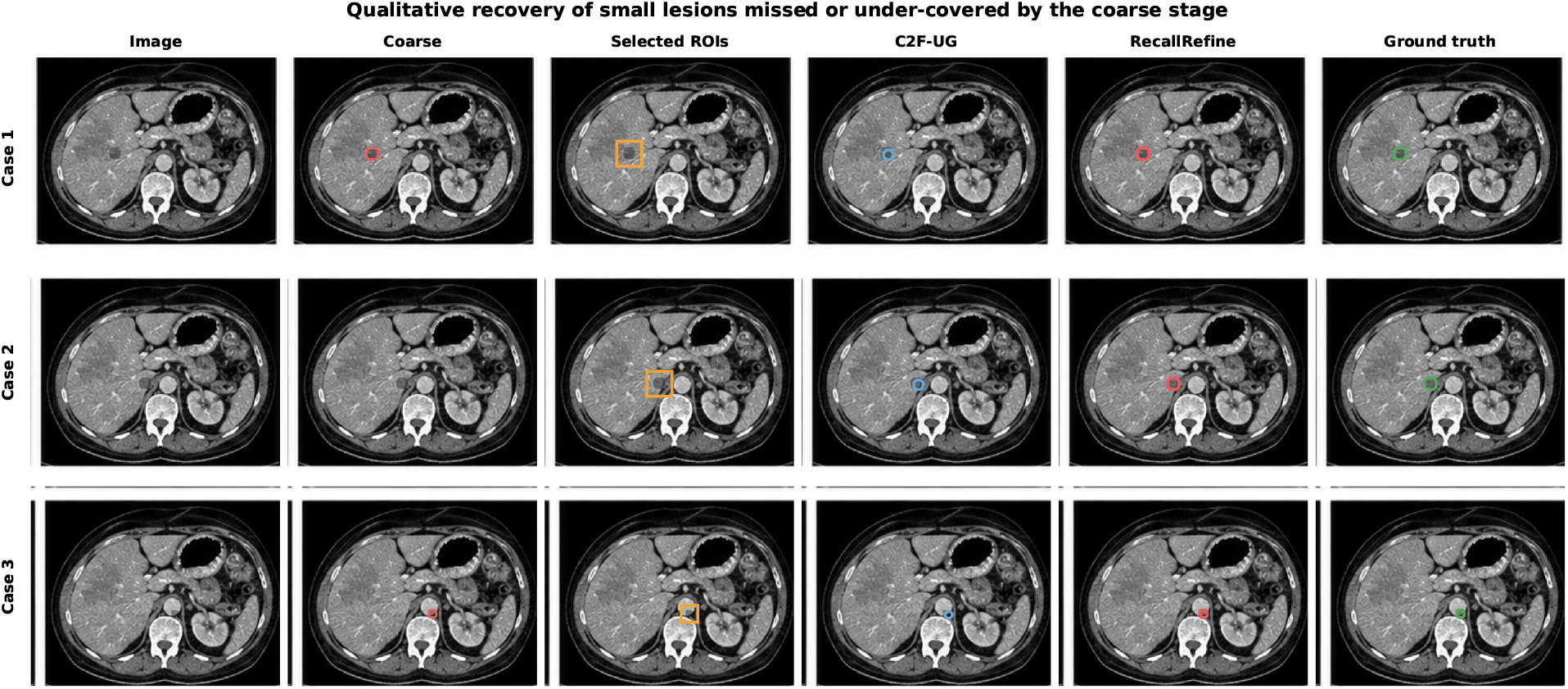
Qualitative examples. Each row shows the input, coarse prediction, selected refinement regions, an uncertainty-guided cascade, RecallRefine, and ground truth. The selected regions explicitly expose whether a lesion was available to the second stage before refinement.

### 4.7. Compute–coverage trade-off

The compute curve in Figure 2c shows that candidate selection changes the accuracy–efficiency frontier. Exhaustive or dense high-resolution models spend computation on large negative regions, whereas a coarse-mask crop is efficient but recall-limited. RecallRefine improves Dice at 148 GFLOPs because the selector allocates a small number of native-resolution blocks to the lesion instances most likely to be discarded by a standard cascade.

## 5. Discussion

The central design choice in RecallRefine is to treat lesion coverage as a first-class learning objective. This is different from simply weighting small lesions more heavily in the segmentation loss. A weighted loss can improve the coarse network, but a difficult lesion may still disappear after thresholding. Coverage supervision instead acts on the *set of regions made available to the fine network*, protecting recoverability even when the initial mask is imperfect.

The method is also distinct from uncertainty-only refinement. Uncertainty is valuable when the model recognizes ambiguity, but modern networks can be confidently wrong.

Feature novelty and component-wise coverage supply complementary signals. Conversely, novelty alone can overselect unusual background. Joint training lets the score network learn how these cues interact under an explicit refinement budget.

There are limitations. First, the component-wise coverage objective requires instance decomposition of the training mask; this is straightforward for disconnected lesions but less well-defined when neighboring lesions merge. Second, the current implementation uses axis-aligned blocks, which are simple and efficient but may be suboptimal for elongated abnormalities. Third, coverage is optimized on the source training distribution; external shifts can still change the relation between candidate scores and lesion presence. These directions motivate future work on shape-adaptive candidates and calibrated coverage under distribution shift.

## 6. Conclusion

We presented RecallRefine, a coverage-constrained coarse-to-fine framework for small 3D lesion segmentation. The method changes the role of the first stage from merely producing a rough mask to constructing a refinement set that preserves difficult lesions. A differentiable component-wise coverage loss, multi-cue candidate scoring, budgeted selection, and context-aware native-resolution refinement together reduce irreversible coarse-stage misses. Across four lesion benchmarks, the largest improvements occur for lesions below 10 mm, supporting the view that *what is routed to the fine stage* is a central learning problem in cascaded medical segmentation.

### Impact Statement

Automated lesion segmentation can support quantitative imaging workflows but should not be interpreted as a standalone diagnosis. The proposed method focuses on computational allocation and lesion coverage; clinical deployment would require external validation, calibration, and workflow-specific review. Because improved sensitivity can increase false-positive burden, both lesion-level recall and false positives per volume should be reported in practical evaluations.

## A. Additional Implementation Details

The candidate generator uses three isotropic side lengths (32, 48, and 64 voxels) with 50% overlap. Coarse probabilities, entropy, and pooled feature novelty are normalized per volume before being passed to the score network. Feature novelty is measured relative to an exponential-moving-average prototype computed from voxels confidently classified as background. The context ring expands the selected ROI by 25% on each side and is encoded separately before fusion.

## B. Candidate Coverage Protocol

We report a lesion as covered when at least 80% of its voxels lie inside the union of selected blocks. We additionally report a relaxed “touched” metric requiring only one overlapping selected block, but use the stricter coverage measure in the main text. For lesions smaller than one candidate block, the distinction is minor; for larger irregular lesions, the stricter metric prevents a selector from receiving credit for only touching a small lesion boundary.

## C. Additional Baseline Details

All dense baselines are trained with their published preprocessing conventions and the same train/validation split. For the matched two-stage baselines, the coarse network, candidate budget, native-resolution refiner, augmentation, and optimizer are held fixed; only the candidate-selection rule changes. This isolates the effect of coverage-constrained selection from architectural differences in the segmentation backbone.

## D. Additional Results

At a 10% candidate budget, RecallRefine preserves 84% lesion coverage, compared with 70% for uncertainty-only selection. At 40%, all selectors approach saturation, reducing the advantage of learned coverage. This is expected: the contribution is most useful when native-resolution refinement is scarce enough that candidate choice matters.

## References

Abraham, N. and Khan, N. M. A novel focal tversky loss function with improved attention u-net for lesion segmentation. In ISBI, 2019.

Antonelli, M., Reinke, A., Bakas, S., et al. The medical segmentation decathlon. Nature Communications, 13: 4128, 2022.

Bilic, P., Christ, P., Li, H., et al. The liver tumor segmentation benchmark (lits). Medical Image Analysis, 84: 102680, 2023.

Hatamizadeh, A., Nath, V., Tang, Y., Yang, D., Roth, H. R., and Xu, D. Swin unetr: Swin transformers for semantic segmentation of brain tumors in mri images. In MICCAI BrainLes Workshop, 2022a.

Hatamizadeh, A., Tang, Y., Nath, V., Yang, D., Myronenko, A., Landman, B., Roth, H. R., and Xu, D. Unetr: Transformers for 3d medical image segmentation. In WACV, pp. 574–584, 2022b.

Heller, N., Isensee, F., Maier-Hein, K. H., et al. The kits19 challenge data: 300 kidney tumor cases with clinical context, ct semantic segmentations, and surgical outcomes. Medical Image Analysis, 67:101821, 2021.

Hu, Y. et al. Uncertainty-aware refinement framework for ovarian tumor segmentation in cect volume. Medical Physics, 2024.

Isensee, F., Jaeger, P. F., Kohl, S. A. A., Petersen, J., and Maier-Hein, K. H. nnu-net: a self-configuring method for deep learning-based biomedical image segmentation. Nature Methods, 18(2):203–211, 2021.

Isler, I. S., Mohaisen, D., Lisle, C., Turgut, D., and Bagci, U. Uncertainty-guided coarse-to-fine tumor segmentation with anatomy-aware post-processing. arXiv preprint arXiv:2504.12215, 2025.

Kervadec, H., Bouchtiba, J., Desrosiers, C., Granger, E., Dolz, J., and Ben Ayed, I. Boundary loss for highly unbalanced segmentation. In MIDL, pp. 285–296, 2019.

Litjens, G., Kooi, T., Bejnordi, B. E., et al. A survey on deep learning in medical image analysis. Medical Image Analysis, 42:60–88, 2017.

Ma, J., Li, F., and Wang, B. U-mamba: Enhancing longrange dependency for biomedical image segmentation. arXiv preprint arXiv:2401.04722, 2024.

Milletari, F., Navab, N., and Ahmadi, S.-A. V-net: Fully convolutional neural networks for volumetric medical image segmentation. 3DV, pp. 565–571, 2016.

Ronneberger, O., Fischer, P., and Brox, T. U-net: Convolutional networks for biomedical image segmentation. MICCAI, pp. 234–241, 2015.

Roy, S., Koehler, G., Ulrich, C., Baumgartner, M., Petersen, J., Isensee, F., Jaeger, P. F., and Maier-Hein, K. H. Mednext: Transformer-driven scaling of convnets for medical image segmentation. In MICCAI, pp. 405–415, 2023.

Sudre, C. H., Li, W., Vercauteren, T., Ourselin, S., and Cardoso, M. J. Generalised dice overlap as a deep learning loss function for highly unbalanced segmentations. DLMIA, pp. 240–248, 2017.

Wang, G., Li, Y., Chen, W., Ding, M., Cheah, W. P., Qu, R., and Shen, L. S3-mamba: Small-size-sensitive mamba for lesion segmentation. In AAAI, volume 39, pp. 7655–7664, 2025.

Xi, S., Wang, S., Safari, M., Hu, M., Tian, Z., and Yang, X. Uncertainty as a foundation for trustworthy medical imaging ai: A comprehensive review. 2025.

Xi, S., Hu, S., Wang, S., Ding, H., Li, Y., He, W., Zhang, K., Del Balzo, L., Zhong, C., Hu, M., et al. Grounding radiology report findings into medical image segmentation. npj Digital Medicine, 2026a.

Xi, S., Hu, S., Wang, S., Safari, M., del Balzo, L., Karim, E. U., Hu, M., Zhang, K., Wang, T., Weichselbaum, R. R., et al. A radiographic world model for clinical reasoning and evidence generation. arXiv preprint arXiv:2609.07719, 2026b.

